# Predicting Phenotypic Traits Using a Massive RNA-seq Dataset

**DOI:** 10.1101/2023.12.05.570195

**Authors:** John Anthony Hadish, Loren A. Honaas, Stephen Patrick Ficklin

## Abstract

Transcriptomic data can be used to predict environmentally impacted phenotypic traits. This type of prediction is particularly useful for monitoring difficult-to-measure phenotypic traits and has become increasingly popular for monitoring high-value agricultural crops and in precision medicine. Despite this increase in popularity, little research has been done on how many samples are required for these models to be accurate, and which normalization should be used. Here we create a massive RNA-seq dataset from publicly available *Arabidopsis thaliana* data with corresponding measurements for age and tissue type. We use this dataset to determine how many samples are required for accurate model prediction and which normalization method is required. We find that Median Ratios Normalization significantly increases performance when predicting age. We also find that in the case of our dataset, only a few hundred samples are required to predict tissue types, and only a few thousand samples are necessary to accurately predict age. Researchers should consider these results when choosing the number of samples in a transcriptomic experiment and during data-processing.

**Author Summary:** Large datasets have become ubiquitous in both research and industry, with thousands and sometimes millions of samples being collected for a single project. In biology a prominent new technology is RNA-seq, which can be used to measure the expression level of thousands of genes for a single sample. These measurements are used for a variety of downstream applications, including predicting phenotypic traits (i.e. height, disease, etc.). A number of experiments have attempted to use RNA-seq data to make phenotype predictions with varying success. This is partially due to the small sample size of their experiments. RNA-seq datasets are currently relatively small--only a dozen to a few hundred samples--due to the cost per sample. This is expected to change as the cost of sequencing decreases. In this paper we create a massive conglomerate RNA-seq dataset from publicly available *Arabidopsis thaliana* RNA-seq data. We use this dataset to determine how many samples are required to accurately predict plant age and tissue type using machine learning models. We also explore the best way to normalize large datasets. Our results show the potential of massive RNA-seq datasets, and can be used to inform experimental design for phenotype prediction.

## Background

Predictive modeling of phenotypic traits using transcriptomic data is a method that is increasing in popularity as the cost of sequencing the transcriptomic decreases. This type of modeling uses count data derived from RNA-seq experiments to identify a set of transcriptomic biomarkers (the expression levels of a set of genes) that are predictive of a phenotypic trait. This type of modeling is feasible because the transcriptome is the most basal phenotype of an organism and is often predictive of both current and future downstream phenotypes. Whereas genetic markers remain constant for the life of an organism, the transcriptome changes readily in response to their environment and to the internal status of an organism. A transcriptomic biomarker is one or more genes whose expression level is indicative of a current or future phenotypic outcome. This means that transcriptomic biomarkers can be used for monitoring and prediction of difficult-to-measure phenotypic traits. Transcriptomic biomarkers are used in medical research with an emphasis on predicting cancer type and stage [1–4]. However, recent research has branched into high-value agricultural crops, where researchers are interested in predicting traits such as flowering time [5], flesh quality traits in apples, pears, and potatoes [6–9], and apple maturity [10].

Despite the increasing interest in transcriptomic modeling of traits, there has not been an investigation of how many samples are required to perform these models. The majority of datasets used for this type of modeling often incorporate only a few dozen to a few hundred samples. This contrasts with predictive models in non-biological research areas which can sometimes have thousands to millions of samples [11,12]. Additionally, RNA-seq datasets have a much higher dimensionality in terms of the number of features (genes) they have when compared to other datasets. An RNA-seq dataset can have measurements for thousands of genes, whereas datasets in other domains typically only have a few hundred [13]. These RNA-seq datasets are referred to as “wide”, containing many features (genes) and relatively few samples. This can present issues for model methods that were created with the intent of only a few features [13,14].

Despite increased interest in biomarker discovery from gene expression, more information is needed to address the question of the number of samples that might be needed to accurately find biomarkers from an underdetermined system using gene expression data. Here we report a study to, first, explore how many samples may be required for transcriptomic modeling of both categorical and continuous phenotypic traits, and second, to identify the effect that RNA-seq count normalization methods have on large disparate RNA-seq datasets. Normalization is a potential acute problem for both large data sets collected over several years or by multiple collaborating groups as well as for large conglomerate datasets with potentially hundreds of different experiments.

In this paper, we create a massive RNA-seq dataset from *Arabidopsis thaliana* (Arabidopsis) retrieved from the National Center for Biotechnology Information’s (NCBI) Sequence Read Archive (SRA) database [15]. We also create a large phenotype annotation dataset from companion data available on NCBI BioProjects database [16,17] which we manually curate. We chose to use Arabidopsis, as it is a model organism in plant science [18] with well-defined physiological stages [19] and has a large amount of RNA-seq data available for it. This makes Arabidopsis an ideal candidate for investigating the size of the dataset required for creating accurate models. The diverse phenotype annotations available for this Arabidopsis data allow us to investigate models for both classification-based variables (tissue type) and continuous variables (age).

We use random forest models [20] for investigating these datasets, as it is robust to outliers and is capable of dealing with a large number of features (in our case genes) [21], and has been shown to be superior to deep learning methods for count data [3]. We also make use of Boruta [22] for feature reduction, and Synthetic Minority Over-sampling TEchnique (SMOTE) [23] for over-sampling (supplementing) sparse data. Additionally, we investigate several normalization methods (Trimmed Mean of M values (TMM) [24], Median Ratios Normalization (MRN) [25], Transcripts Per kilobase Million (TPM), and No Normalization (NoNo) to determine which is best for dealing with large conglomerate datasets.

The following research seeks to address how many samples are required for transcriptomic modeling of both categorical and continuous phenotypic traits, identify the best normalization method for dealing with large conglomerate RNA-seq datasets, and demonstrate how large conglomerate datasets can be mined for additional information beyond their initial intent. This has relevance for experimental design interested in identifying transcriptomic biomarkers for both categorical and continuous phenotypic traits. We found that MRN normalization performed better than other forms of normalization when predicting age, whereas normalization had no impact when predicting tissue type. We also found that a few hundred samples are sufficient for predicting our categorical variable of tissue type, whereas a few thousand samples are required for modeling the continuous variable of age.

## Data Description

### RNA-seq Data Pre-Processing

Arabidopsis RNA-seq data was retrieved from the National Center for Biotechnology Information (NCBI) Sequence Read Archive (SRA) [15] using the following search parameters: txid3702[Organism:noexp] AND (“biomol rna”[Properties] AND “platform illumina”[Properties] NOT “strategy wxs”[Properties] NOT “strategy targeted capture”[Properties] NOT (“strategy other”[Properties] NOT (“library selection pcr”[Properties] NOT “library selection padlock probes capture method”[Properties] NOT “library selection hybrid selection”[Properties] NOT “library selection other”[Properties]))) NOT (“strategy chip”[Properties] NOT “strategy mre seq”[Properties] NOT “strategy atac seq”[Properties] NOT “strategy faire seq”[Properties] NOT “strategy mnase seq”[Properties] NOT “strategy dnase hypersensitivity”[Properties] NOT “strategy medip seq”[Properties] NOT “strategy mbd seq”[Properties] NOT “strategy bisulfite seq”[Properties]) NOT “filetype bam”[Properties] AND (“biomol rna”[Properties] AND “platform illumina”[Properties] AND “filetype fastq”[Properties])

This search string retrieves all *Arabidopsis thaliana* (Arabidopsis)(NCBI txid3702) RNA-seq data in the fastq format created using an Illumina sequencing machine (Illumina, San Diego, CA, US). A total of 74833 SRR (RNA-seq runs) were identified with these parameters corresponding to 60062 SRX (RNA-seq experiments). An SRX experiment can consist of multiple RNA-seq run files.

The list of 74833 SRR accessions was used with the GEMmaker workflow [26] automatically retrieves from NCBI and processes the SRR files. For these data, GEMmaker was run using Kallisto [27] for expression quantification. For genome alignment, GEMmaker was given the Arabidopsis genome (TAIR 10 assembly, Araport 11 annotations) retrieved from The Arabidopsis Information Resource (TAIR) [28]. Data was split into batches of ∼5000 SRR numbers for processing. While splitting the data is not necessary for GEMmaker, doing so allowed for execution on multiple queues on Washington State University’s high-performance computing cluster “Kamiak” and took approximately 3 months to complete using available nodes at the time. During execution, 5515 SRX experiments were removed due to improper SRA file formatting, SRA file corruption, or empty SRA files (9.2% removed). The resulting Gene Expression Matrix (GEM) consisted of 54547 samples and 48359 genes representing 2605 NCBI BioProjects. After the creation of the GEM, custom Python code was used to remove samples that did not have at least 1 million reads and did not have at least 70% of reads aligning with the Arabidopsis genome. A total of 288 sample pairs were noticed to be identical, and on closer inspection, it was determined that the same RNA-seq sample had been uploaded to NCBI multiple times (at least twice, as many as five times) under different names (and often different phenotype annotations). These were removed from consideration. Within the remaining samples, genes with expression in less than 1000 samples over a count of 10 (in the NoNo dataset) were removed. This was an ad hoc filter based on “filterByExpr” function of the edgeR package [24], with a number of different posible combinations of “samples” and “count” variables assessed to see how this would influence gene count (S1 Figure). This filtering resulted in a final GEM consisting of 32044 samples and 43224 genes.

Four separate GEMs were created to test model performance for different normalization methods: Trimmed Mean of M values (TMM), Median Ratios Normalization (MRN), Transcripts Per kilobase Million (TPM), and No Normalization (NoNo). TMM normalization [24,29] and MRN normalization [25] were performed using the Python “conorm” package 1.2.0 [30]. TPM and NoNo normalization values were an output of Kallisto [27]. How these normalizations impacted sample count is visualized as S2 Figure. We have made these GEMs publicly available on Zenodo at the link https://zenodo.org/records/10183151

### Sample Phenotype Annotations Pre-Processing

Sample phenotype annotations were retrieved from the NCBI BioProject database [16,17] using BioSampleParser which was slightly modified to check for successful data retrieval [31]. Phenotype annotations were retrieved for 48696 NCBI BioSamples, representing data from 2643 BioProjects. A total of 668 different phenotype annotation classes (e.g., “tissue”, “age”, “organism”, “title”, etc.) were retrieved. Most of these phenotype annotation classes were present for one or a few BioProjects, and therefore were sparse, only containing a few samples (S3 Figure). These sparse classes were ignored. Phenotype annotation classes assigned to over 5000 RNA-seq samples are visualized in Figure 1 A. For this experiment, we used the phenotype annotation classes “tissue” and “days” because of their large number of associated RNA-seq samples and biological relevance. Additionally, “Tissue” is categorical and can therefore be modeled as a classification problem (i.e., which tissue did a sample come from), whereas “days” is continuous and can be modeled as a regression problem (i.e., how old is this sample).

**Figure 1:**
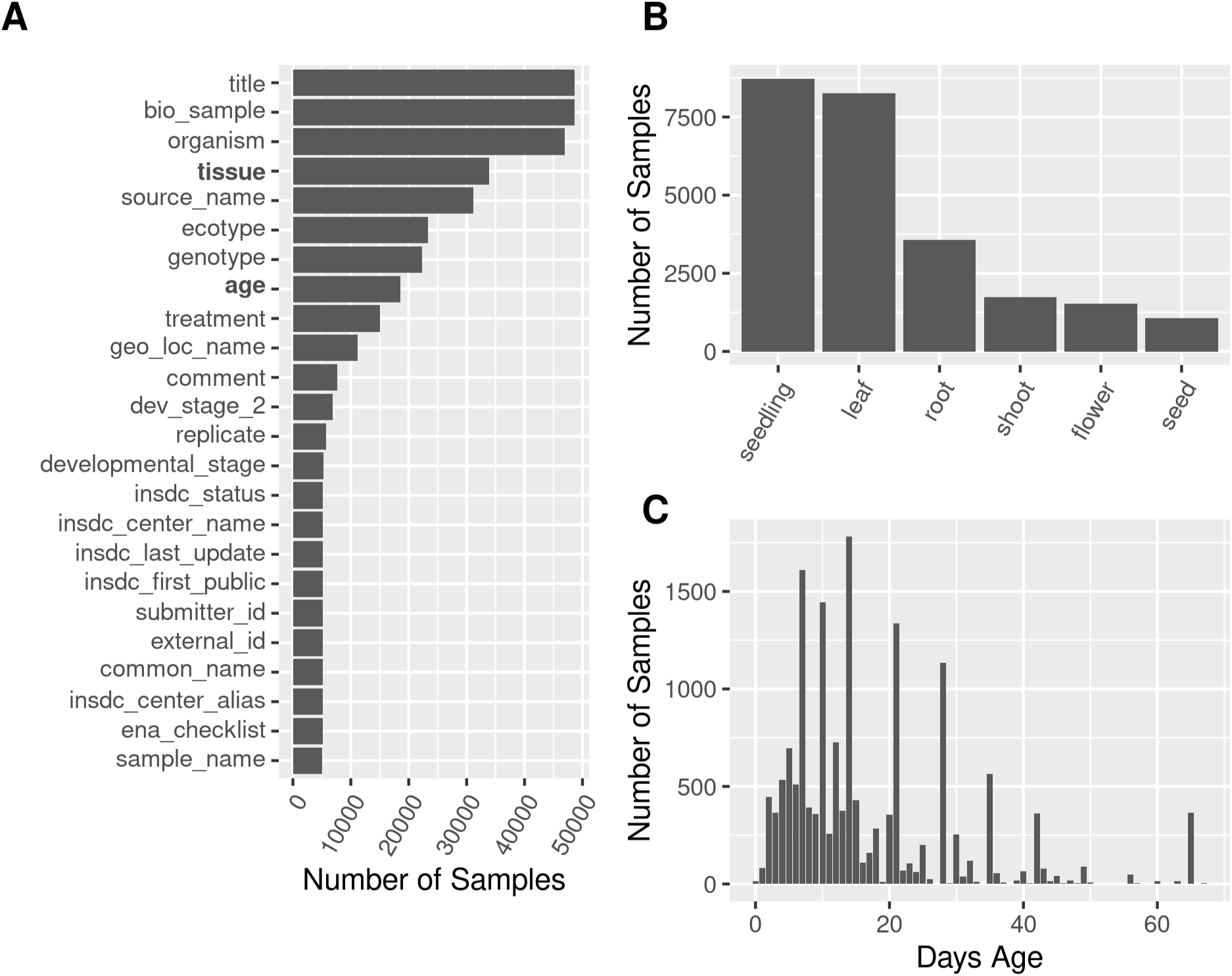
Phenotype annotations retrieved from NCBI for the Arabidopsis dataset. **A)** Available phenotype annotation columns. The y-axis represents the name of the phenotype annotation column, and the x-axis represents the number of samples that have that phenotype annotation column. Phenotype annotation columns can be both biological related and technical related. Phenotype annotation columns with at least 5000 samples are shown. For this experiment, we decided to concentrate on tissue and age. **B)** Tissue annotations. The x-axis represents the tissue annotation type, and the y-axis represents how many samples have that annotation. **C)** Age annotations. The x-axis represents how old the sample annotation is (rounded to the nearest day), and the y-axis represents how many samples are that age. Age annotation was reported on NCBI in many ways, with a breakdown figure reported as (S4 Figure).

The “tissue” phenotype annotations from NCBI included a total of 1186 unique annotation terms (Figure 1 B) for the RNA-seq samples. Not all samples had “tissue” phenotype annotations, and some annotations were not useable. Due to inconsistent use of terms such as misspellings and ambiguity, these terms required manual curation. In summary, we formed six tissue categories with the following terms: “leaf”, “seedling”, “shoot”, “seed”, “root”, and “flower” (referred to as the “tissue-6” dataset). In addition, a second dataset consisting only of “leaf”, “seed”, “root”, and “flower” was created to test on precise labels, and is referred to as “tissue-4”. As part of our curation process, we made several changes. First, misspellings were corrected. Generic terms (e.g. “the plant”, “whole” “col-0”), tissue types created for specific laboratory applications which are unlike their donor tissue (e.g., “protoplasts”, “in vitro cotyledon”, “tissue culture callus”), and unknown or inappropriate values (e.g., “usa”, “p100”, “liver”) were excluded. A notable conglomeration of samples combined “inflorescence” (e.g., “immature inflorescence”, “plant inflorescence”, “inflorescence containing stage 8 and younger flowe”) terms with “flower” related terms (e.g., “mature flower”, “immature flower bud cluster”, “young_flower_control”). Our “seedling” term was defined as plants younger than 6 days post germination (stage 1, before first primary leaves, plate-based) [19], but it should be noted that many samples labeled with the term “seedling” had no information about the day they were collected and were still included. Terms related to “rosette” (e.g., “complete rosette”, “aerial rosette tissue”, “entire vegetative rosette”) were manually changed to “leaf” unless other information was included (e.g., “rosette and inflorescence”). The “shoot” term was assigned to a large group of samples (1142 samples) but is ambiguous in its meaning, as the flat, rosette nature of adult Arabidopsis plants means that this tissue type is difficult to define in plants over a few days old. After manual correction, the tissue labels were combined with the filtered GEMs, resulting in a data frame of 16271 RNA-Seq samples from 1128 BioProjects across these 6 tissue types (Table 1).

**Table 1:** Distribution of the different tissue categories. BioProjects with multiple tissue categories are included in all counts.

| Tissue Category | Number of Samples | Number of BioProjects |
| --- | --- | --- |
| flower | 723 | 79 |
| leaf | 5321 | 364 |
| root | 2436 | 202 |
| seed | 689 | 59 |
| seedling | 5960 | 440 |
| shoot | 1142 | 79 |

The age annotation consisted of 857 unique values across all samples. Like tissue, age was reported by researchers in multiple ways and not all samples reported age. Age was reported from the time of “seeding”/”germination”, after an event such as “flowering” or “inoculation”, or as a raw number without any additional information. Age was also reported with different terms such as “day”, “week” and “month”. Some age values were improper (“Austria: Innsbruck”, “Nitrogen, plus Cycloheximide, Dexamethasone”, “environmental-water”, etc.), were not day specific (“just prior to or at bolting”, “Adult”, etc.) or implausible (“6month”, “67 years”, etc.). Such values were excluded. A summary of the valid age annotation values is available in Table 2 and visualized in S4 Figure. For our experiment, we used age values reported with the term “Days” in the training and testing of models and used other age values to verify the models. Those RNA-seq samples with age in “Days” values combined with the cleaned phenotype annotations are hereafter referred to as “Days” Age annotation Labeled (DAL) dataset. DAL is a subset of the age dataset. Additionally, we used the time period of 0 to 30 days and excluded later days because of sparsity after 30 days (Figure 1 C) preliminary models tended to perform poorly when they were included. RNA-seq samples with ages between 0 and 30 were combined with the filtered GEMs, which resulted in a dataset with 6136 samples from 485 BioProjects.

**Table 2:** Distribution of the different age categories.

| Age Category | Number of Samples | Number of BioProjects |
| --- | --- | --- |
| Reported as Time |  |  |
| **Days (DAL)** | 6457 | 508 |
| **Weeks** | 2263 | 195 |
| **Months** | 38 | 5 |
| **Number Only** | 1177 | 67 |
| After Event |  |  |
| **Days After Sowing (DAS)** | 514 | 19 |
| **Days After Germination (DAG)** | 445 | 45 |
| **Days After Pollination** | 112 | 7 |
| **Days After Flowering** | 68 | 3 |

After combining the GEMs with the six tissue categories or age in days, respectively, the resulting data frames were split into training and testing datasets. These splits considered BioProject as a batch effect and required that samples from a single BioProject are all either in the training dataset or the testing dataset. This requirement was meant to prevent the overfitting of models due to latent covariates resulting from sample preparation and sequencing (sequencing depth and sample preparation type) within BioProjects. For the tissue dataset, a total of 12749 samples from 902 BioProjects were in the training dataset and 3522 samples from 226 BioProjects were in the testing dataset (the testing set has 28% of the samples and 25% of the BioProject). S1 Table shows a breakdown of these splits based on the 6 conditions. For the DAL dataset, a total of 5062 samples from 388 BioProjects were in the training dataset and 1074 samples from 97 BioProjects in the testing dataset (the testing set has 21% of the samples and 25% of the BioProjects). These splits were used for all models except if otherwise stated.

### Model Parameter Optimization

Random forest model parameter optimization was carried out using the RandomizedSearchCV class from the Python package Scikit-learn (sklearn) [32]. Random Forest models were created using the RandomForestClassifier class for classification of tissues and the RandomForestRegression class was used for age prediction. In both cases, the parameter search space iterated over the following grid: ‘bootstrap’: [True, False], ‘n_estimators’: [100, 300, 500, 1000, 1500, 2000], ‘n_estimators’: [3, 5, 10, 20], ‘max_depth’: 3 to 100 at an interval of 3), ‘min_samples_split’: [1,2,4], ‘min_samples_leaf’: [3, 5, 10, 20, 30], ‘max_features’: [’sqrt’, ‘log2’]. These parameters were sampled 100 times for each of the four filtered GEMs (NoNo, and three GEMs normalized by TMM, MRN, and TPM respectively). Five-fold cross-validation was performed in each model instance, with folds accounting for BioProject batch effect in a manner similar to the input dataset. Evaluation of each instance was performed using F1 and Accuracy for the tissue classification problem and r^2^ for the age regression problem to determine the optimal model parameters and compare the effect of normalization methods of the GEMs.

## Analyses

### Assessing Phenotype Annotation Accuracy

The datasets used in this project are conglomerates of many BioProjects, with samples and labels being collected and classified in many ways by different researchers. This creates the possibility that our models are fitting on the variation between projects, and not on the actual biological traits of interest (i.e., overfitting). To test that our RNA-seq samples contained information that was reflected in their assigned labels, we conducted a randomization strategy, where a percentage of the dependent variable (tissue or age) from 0 – 100% was randomized in the training dataset. If the model is fitting on true biological information, then model accuracy should decrease over increased randomization. Model creation was done using the optimized parameters found for each respective dataset, with respective accuracy assessments (F1 and Accuracy for tissue-6 and tissue-4, r^2^ for DAL).

### Model Performance Metrics

Models using all available data for the tissue dataset (six categories) and DAL dataset (from 0-30 days) were assessed and visualized using respective optimized parameters. Visualization for the categorical tissue dataset uses a confusion matrix and visualization of the quantitative DAL dataset compared to predicted and actual data as a scatterplot.

To determine how many samples are required to accurately predict “tissue” and “age” in our datasets, modeling was performed using different size sample sets. This was performed in an iterative manner, starting with a small training dataset, and gradually adding samples. We used two approaches. The first approach added samples from the training dataset by BioProject: each iteration added all the samples from 10 random BioProjects. The average number of samples added at each iteration was 92.9 (std 87.3) for the tissue dataset and 128.2 (std 42.3) for the DAL dataset. For the second approach, the entire training datasets were randomized (irrespective of BioProject) and samples were randomly added in batches (for age: batch size of 20 for the first 500 samples added, batch size of 40 for the next 500 samples added, batch size of 100 for the next 1000 samples, and batch size of 200 for the remaining samples (up to 5062). For tissue-4: batch size of 10 for the first 500, batch size of 30 for the next 500, batch size of 60 for the next 1000, and batch size of 200 for the remaining. For tissue-6: batch size of 20 for the first 1000, batch size of 100 for the remainder, see **S5 Figure** x-axis). In both cases, the same testing dataset was used every time (independent BioProjects). The first approach is intended to evaluate the required number of independent BioProjects for good model performance. The second approach is intended to simulate a single homogenous--albeit high variance--dataset. This is to simulate how many samples a researcher would need to gather if they were interested in replicating this independently. Additionally, both methods were performed on a reduced tissue dataset which excluded the categories “shoots” and “seedlings”. This reduced tissue dataset with only 4 categories (”leaf”, “seed”, “root”, and “flower”) is referred to as the tissue 4 dataset.

### Models Using Germination and Sowing Dates

Within the annotations for age were samples labeled with terms related to “Days After Germination” (DAG) and “Days After Sowing” (DAS). These two terms are more specific than data in the DAL dataset, but there were fewer BioProjects using these labels. The DAG dataset had 530 samples from 52 BioProjects and the DAS dataset had 873 samples from 19 BioProjects. Two new random forest regression models were created using just the data from these respective labels. Testing was done in the same manner as described above with the DAL dataset. Additionally, both the DAG and DAS models were used to predict day within the DAL dataset to determine if they were more accurate at predicting DAL due to their more specific nature.

### Synthetic Data and Balancing

Data for the DAL dataset were supplemented using Synthetic Minority Over-sampling TEchnique (SMOTE) [23]. This was done to increase the amount of data available for days with few samples to see if this increased model performance. Prior to SMOTE, dates with half days (e.g., 5.5 days) were rounded up to the nearest day. Days with less than 10 samples were removed, as SMOTE is inaccurate with small sample sizes (days 0,1,19,27, 29 were removed). SMOTE oversampling was performed using the imbalanced-learn package (version 0.11.0) with default parameters [33]. This resulted in a new data frame that contained 16112 samples, over three times the amount of the original training dataset size of 5062. The distribution of samples before and after SMOTE is visualized in S6 Figure. Synthetic Data was not created for the tissue dataset due to already high model performance.

### Feature Selection and Evaluation

Feature selection was performed on the DAL, tissue-6, and tissue-4 datasets to determine which genes were most important for predicting tissue and days of age. Boruta feature selection was performed on both the DAL and the tissue dataset using their respective optimized parameters. Borutapy v 0.3 [22,34] parameters were set to *n_estimators=*’auto’ (defaults to 1000), *max_iter=*200, *perc* = 100 for both the tissue and DAL datasets. The results of these Boruta runs are sets of genes that can predict the dependent variable (i.e., tissue or days of age) better than a randomized version of themselves. Genes selected by Boruta were used to create new datasets: a new DAL Boruta Dataset which had 15024 genes remaining, a new tissue-6 Boruta Dataset which had 20017 genes remaining, and a new tissue-4 Boruta dataset which had 7837 genes remaining. These reduced Boruta datasets were then used to create new random forest models. While boruta is able to determine if a gene is better than a random version of itself, it does not rank the genes on their importance to the model. To generate feature importance scores, new random forest models were created.

## Results

### Optimizing Input Parameters and Assessing Normalization Method

Normalization methods dramatically impacted GEM values and therefore total reads per sample. A distribution of the number of reads per sample for all 4 normalization methods is visualized as S2 Figure. These normalized GEMs were used to create models for both tissue classification and age prediction. For the tissue classification, no significant difference in model performance was seen between the 4 normalization methods (one-way ANOVA, p = 0.485) (Figure 2 A). In contrast, a significant difference in performance was observed between different normalization methods for the DAL dataset (one-way ANOVA, p = 0.00012), with a posthoc Tukey’s Honest Significant Difference (HSD) test revealing that MRN performed significantly better than either NoNo, TMM, or TPM (Figure 2 B) (S2 Table). While no difference was seen in normalization methods for the tissue classification, it was decided to use MRN for the remaining analyses due to its increased performance with the age model.

**Figure 2:**
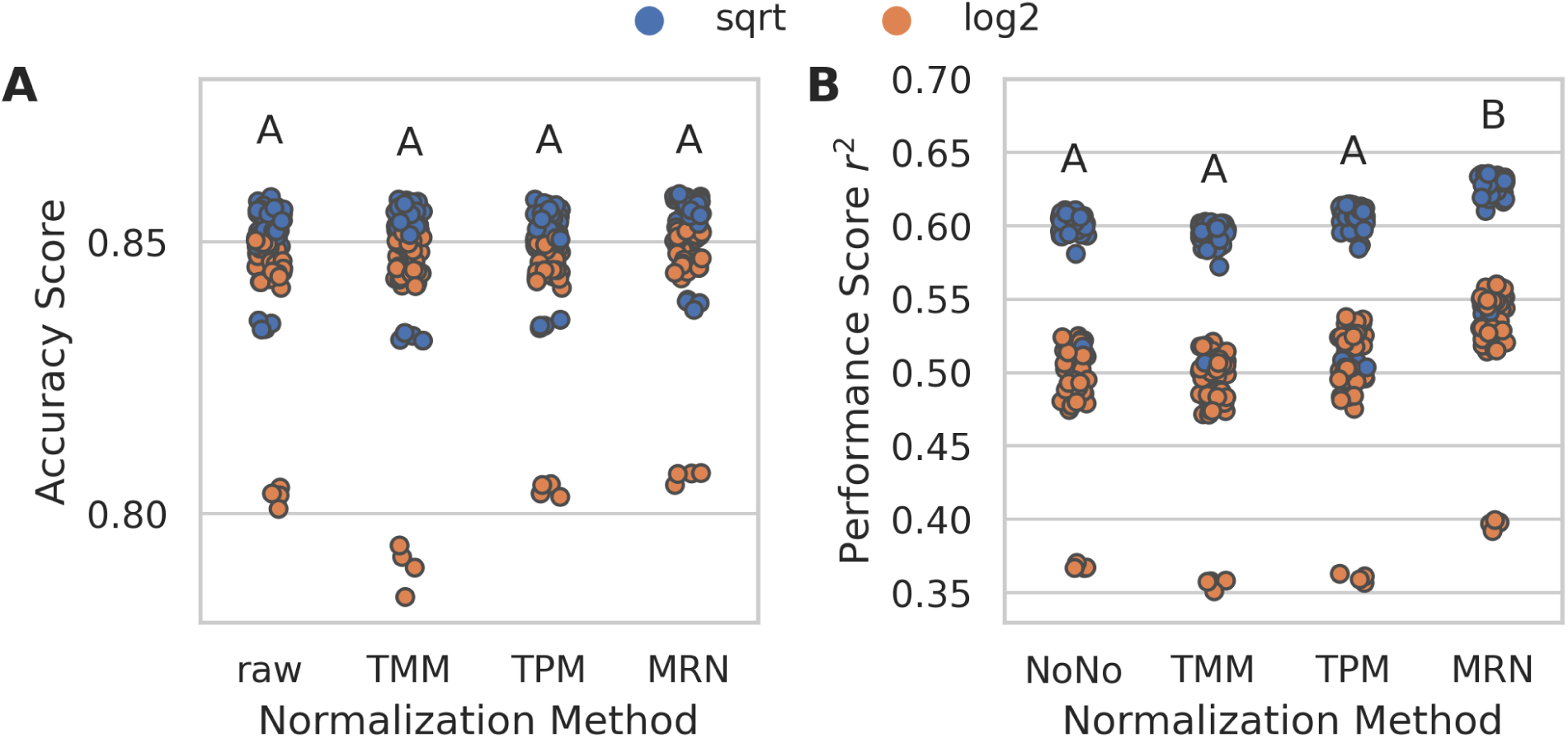
Assessing parameter optimization and normalization methods **A)** Tissue model. There were no significant differences between normalization methods (one-way ANOVA, p = 0.475) **B)** Age model results. A and B represent significant differences based on Tukey’s honestly significant difference (HSD) test S2 Table. Of the 4 normalization methods, MRN had the best performance on average. We also saw separation in the chart, which was most impacted by the parameter “max_features” (colored points). Additional “max_features” values were evaluated and can be seen in S7 Figure. Points that are separated from their respective colored clusters (log2 around 0.36 and sqrt around 0.53) were found to be models where “max_depth” was too low and set to only 3 S8 Figure.

In addition to evaluating different normalization methods, random forest model parameters were tested with different values to identify optimal model performance. Parameter sets for both the DAL and tissue datasets were tested in a similar manner. For tissue classification the optimal parameters were ‘n_estimators’: 1500, ‘min_samples_leaf’: 3, ‘max_features’: ‘sqrt’, ‘max_depth’: 48, ‘bootstrap’: False (S3 Table). For the age model, the optimal parameters were *’n_estimators’*: 1500*, ‘min_samples_leaf’*: 3, *’max_features’*: ‘sqrt’, *’max_depth’*: 48, *’bootstrap’*: False (S4 Table). However, it was found that reducing ‘*n_estimators’* to 300 had negligible impact on the r^2^ score while dramatically reducing runtime (300: mean r^2^ 0.555, std 0.051, 1500: mean r^2^ 0.563 std 0.049, ANOVA: F-Statistic: 1.0875, P-value 0.2987). Therefore, the remaining analyses used parameters *’n_estimators’*: 300*, ‘min_samples_leaf’*: 3, *’max_depth’*: 7, *’bootstrap’* False. Furthermore, it was decided to set *max_depth* at 7 for both models, as this lower value performed nearly as well as higher parameters and reduces the chance of overfitting the model. The dramatic difference in performance for *max_features* for the age dataset (Figure 2 B)—and to a lesser degree in the tissue dataset (Figure 2 A)--warranted additional investigation. A GridSearch optimizing for *’max_features’* revealed optimal *’max_features’* = 700 for tissue (S7 Figure A) and *’max_features’* = 1000 for DAL (S7 Figure B). These parameters consider the tradeoffs between model performance and model speed, which has an impact when training Boruta models in later steps.

### Optimal Parameter Performance

Tissue classification using optimal parameters showed an F1 score of 0.942 and a model accuracy of 0.948 for the training dataset and an F1 score of 0.739 and an accuracy of 0.792 for the testing dataset. Confusion plots demonstrating the results of each classification are visualized in Figure 3, with subplots A and B showing the training and testing data performance, respectively. Values along the diagonal are the number of correctly predicted labels and those non-diagonal values are incorrect predictions.

**Figure 3:**
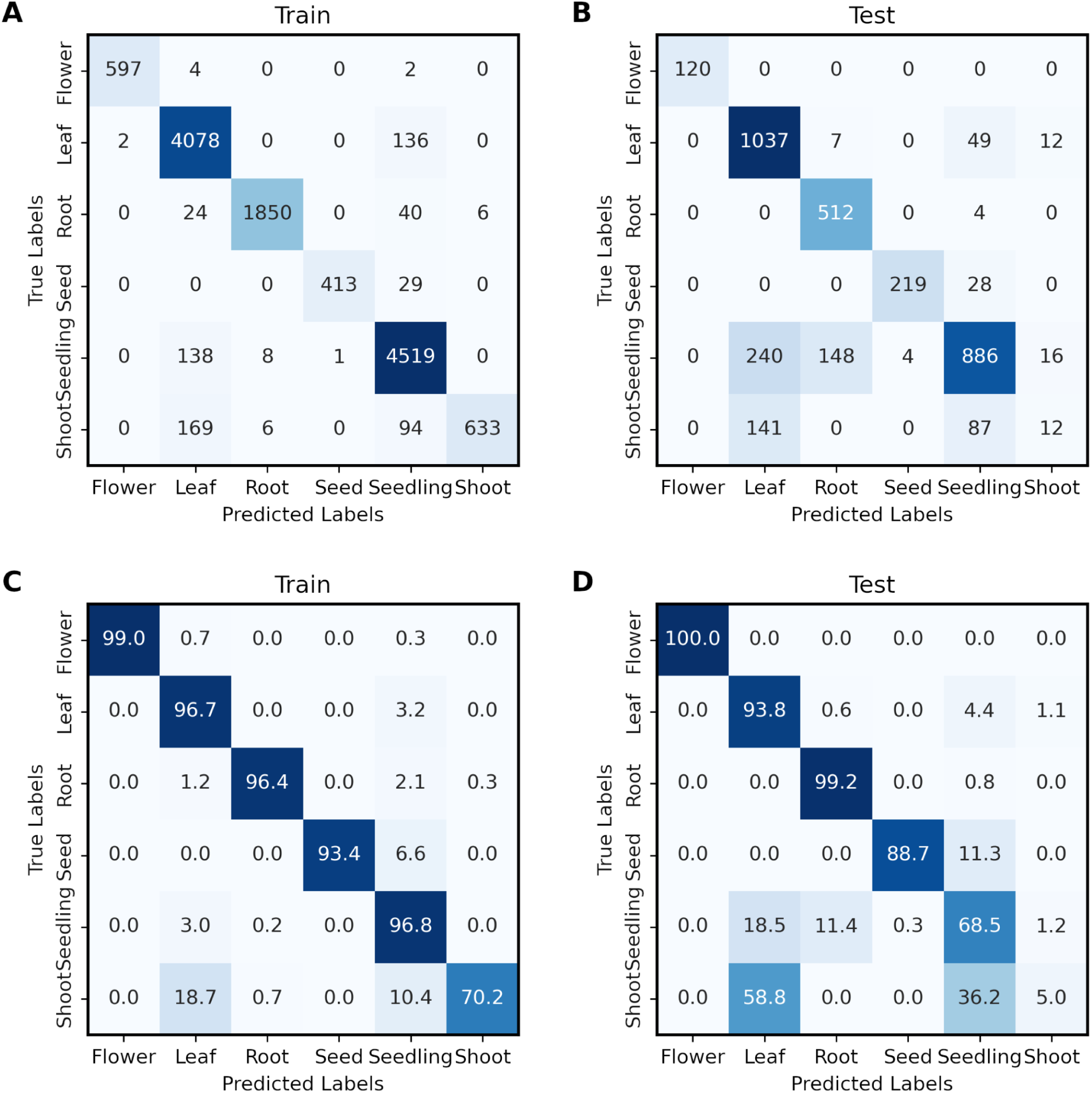
Confusion Matrices of Tissue Data. **A)** Unnormalized (values represent actual count) train **B)** Unnormalized test **C)** Normalized train (values in each row and column sum to 1) **D)** Normalized test.

Subplots C and D show the same results but with counts normalized such that the sums of columns and rows are 1. These confusion matrices show that labels for “flower”, “leaf”, “root” and “seed” performed the best, whereas labels for “seedling” and “shoot” did not perform as well, likely due to their ambiguity, as we saw some annotations labeled as “seedling” which were plants that should have one or more pairs of true leaves and “shoot” is difficult to define except in plants before they develop primary leaves. Therefore, an additional model was created excluding “seedling” and “shoot” (referred to as tissue-4). The tissue-4 model had an F1 score of 0.996 and a model accuracy of 0.995 for the training dataset and an F1 score of 0.995 accuracy 0.994 for testing. A confusion matrix showing the breakdown of categories is shown in Figure 4.

**Figure 4:**
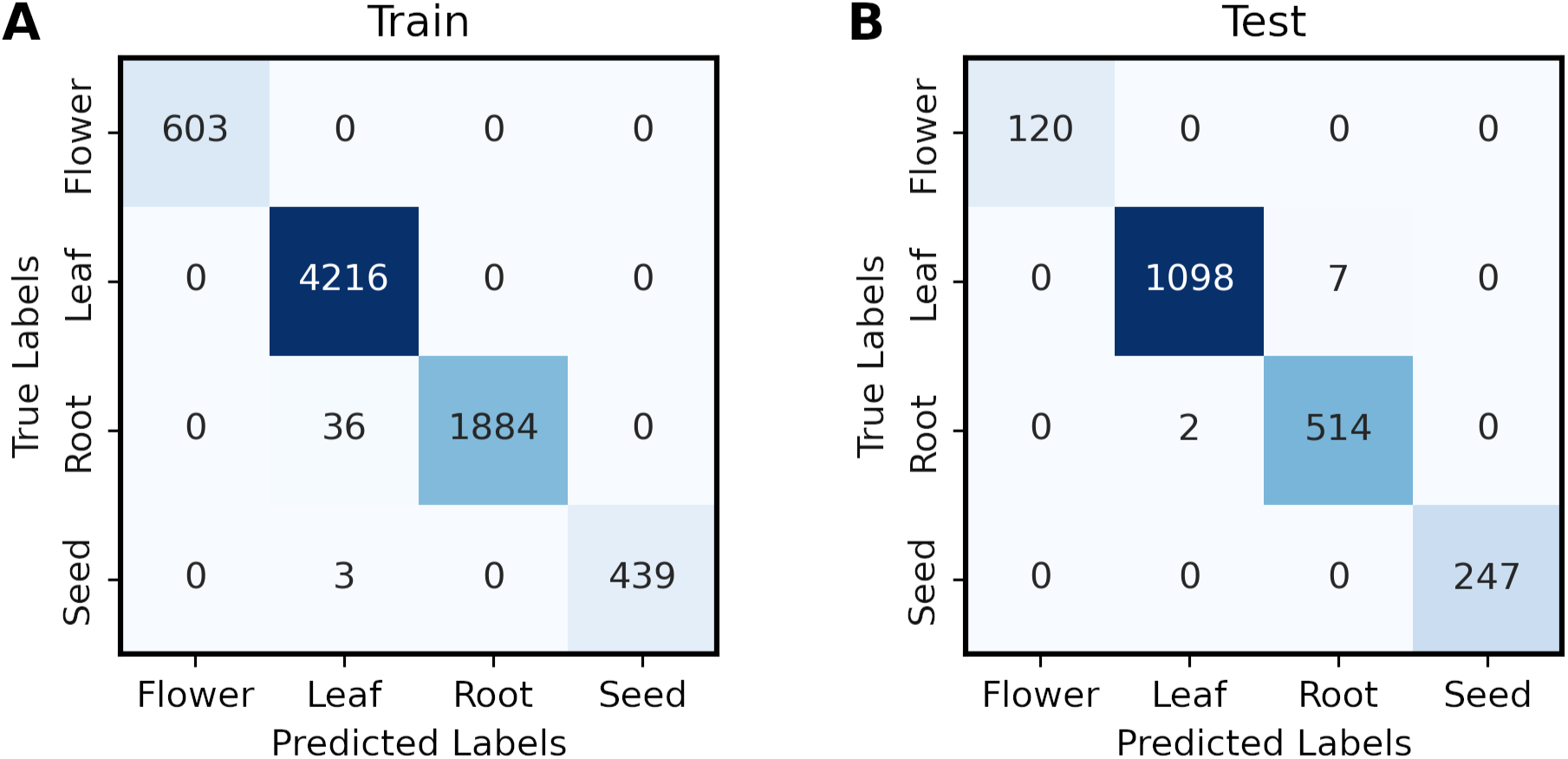
Confusion matrices of tissue-4 models. **A)** Training dataset accuracy. **B)** Testing dataset accuracy.

Age prediction using optional parameters in the regression model for the DAL model had an r^2^ of 0.9409 and Root Mean Square Error (RMSE) of 1.6125 for the training dataset and an r^2^ of 0.5524 and RMSE of 4.4423 for the testing dataset (Figure 5). Two other models were created using data from the complete “day” dataset. The first model was for predicting age from samples that were labeled as Days After Germination (DAG) and the second was for Days After Sowing (DAS). The DAG model had an r^2^ of 0.9983 and RMSE of 0.2992 for the training, and an r^2^ of 0.4493 and RMSE of 0.46904 for the testing. The DAS model had an r^2^ of 0.9995 and RMSE of 0.1231 for the training, and an r^2^ of –0.5712 and RMSE of 3.6729 for the testing. Both the DAG and DAS datasets had substantially fewer samples (distribution visualized in S9 Figure A and D respectively) than the DAL model. The DAS testing model only had testing data for 4 time points which allowed it to achieve an unwarranted better RMSE than the DAL model (which the very poor r^2^ of –0.5712 revealed). We used the above models (tissue-6, tissue-4, and DAL) to predict annotations for the entire RNA-seq dataset, including previously unannotated samples. This table of predictions is available as S5 Table, which also includes information about the train test splits used for each model in this paper.

**Figure 5:**
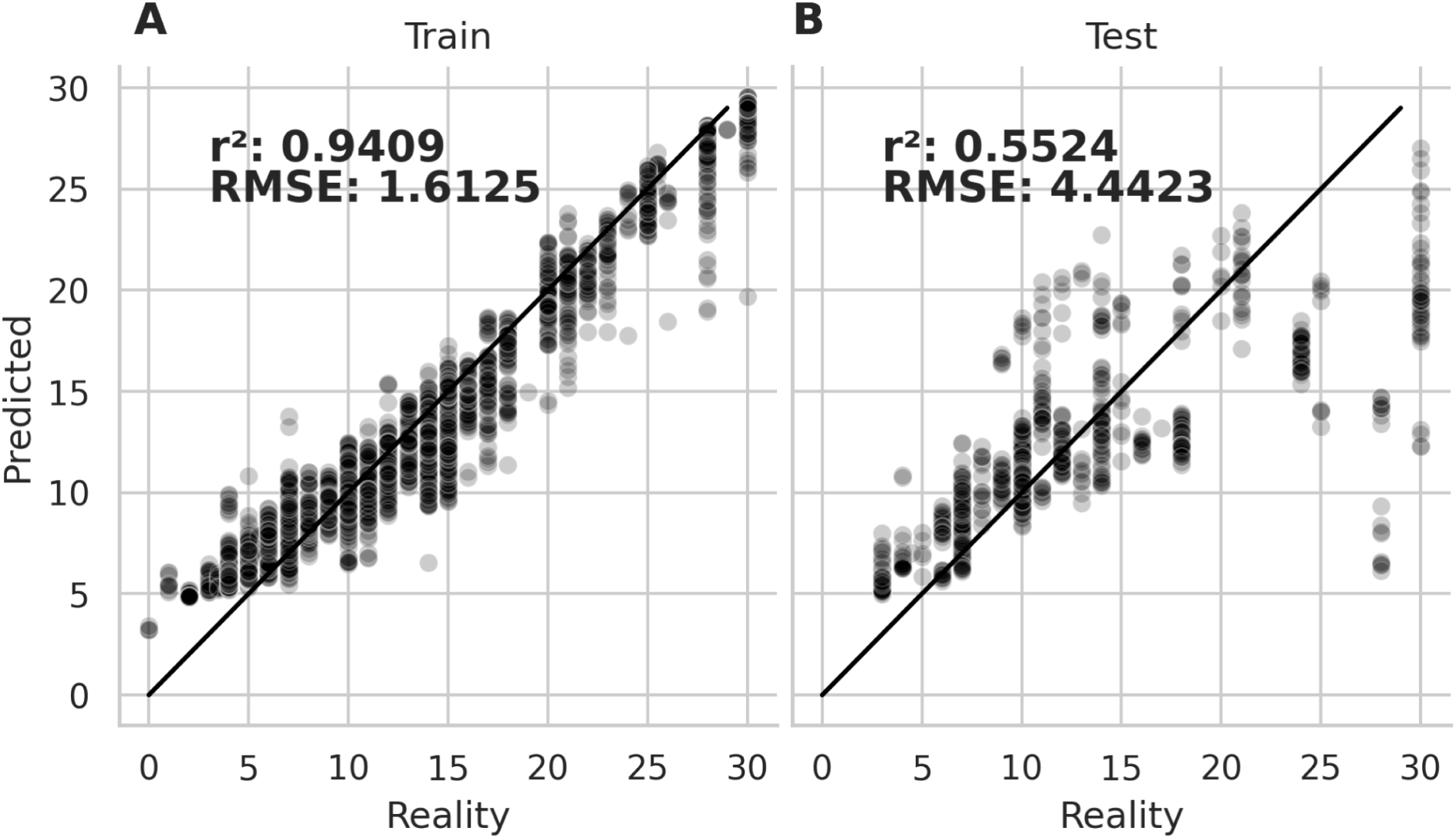
Model performance for DAL model. **A)** Results from the training dataset, and **B)** results from the testing dataset. The x-axis represents the actual age while the y-axis represents the age predicted by the model.

The performance of the DAG and DAS models was also performed using the DAL dataset as testing data (S10 Figure). The model performance of the DAG and DAS models on the DAL dataset was lower than the DAL model. This lower score indicates that the DAG and DAS datasets may be overfit due to the limited data.

An additional model was created using the DAL dataset supplemented using SMOTE. SMOTE synthesizes new data from minority classes (i.e., time points with fewer samples). It does this by drawing lines between random samples of closely related features of the minority class in the feature space. A random point along this line is taken which results in a new synthetic sample that has a resemblance to the two parent samples [23]. SMOTE is effective because it creates new data points which have a plausible feature space as compared to the class they were created from [23]. The SMOTE over-sampled DAL dataset contained 16112 samples, over three times the amount of the original training dataset size of 5062 (S6 Figure). The DAL SMOTE model offered a modest improvement over the DAL model, with an r^2^ of 0.977 and RMSE of 1.2335 for the training dataset and an r^2^ of 0.563 and RMSE of 4.3896 for the testing dataset (S11 Figure).

### Assessing Phenotype Annotation Accuracy

Phenotype annotation accuracy was assessed using randomization. For both tissue classification and DAL prediction, between 0 and 100% of the tissue labels or age values for the training datasets were randomly shuffled. These randomized training datasets were used to create a model, which was then evaluated on the testing dataset. Both models saw decreases in the performance for both training and testing with increasing randomization (Figure 6). The tissue classification did not lose substantial performance until higher amounts of randomization were introduced (Figure 6 A). The DAL model showed decreasing performance over increasing randomization, reaching nearly an r^2^ of 0 at 100% randomization (Figure 6 B).

**Figure 6:**
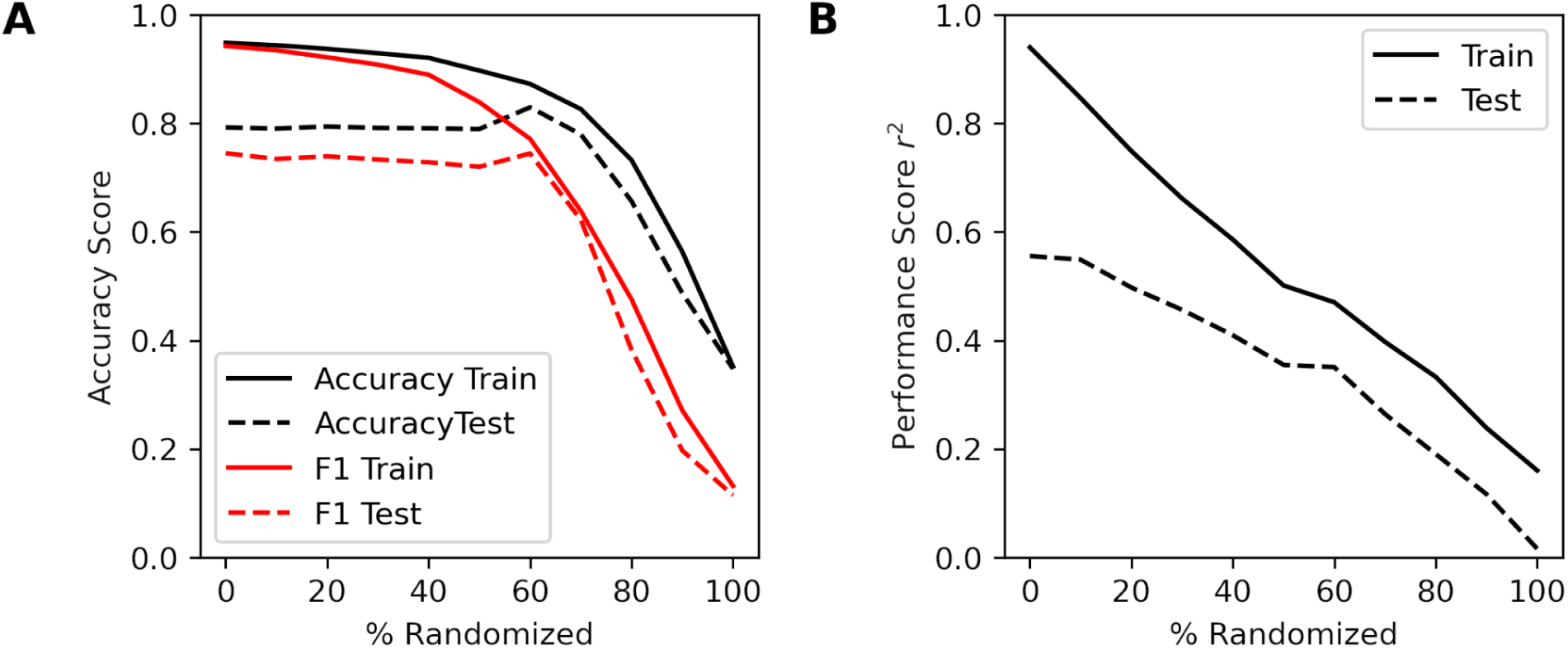
Model accuracy over an increasing amount of randomization for **A)** tissue, measured using both accuracy and F1 score, and **B)** age, measured using r^2^. As the percent of samples with randomized tissue labels or age values increased, accuracy in the model both with the training and testing datasets decreased.

### Number of Samples Required

The number of samples required for maximum accuracy was assessed for both models. Samples were randomized using two different approaches: either according to BioProject or randomly from the training datasets. The intent of this experiment was to determine the number of samples required to reach a plateau of model performance.

Performance results from the model where samples were added randomly from the training datasets for the tissue-4 classification models are shown in Figure 7 A for a F1 range from 0.850 to 1 (for the full range see S12 Figure A). For this randomization approach, the model’s accuracy was at near maximum after adding only a few hundred samples. This is in contrast with the regression DAL model, which reached its maximum only after a few thousand samples Figure 7 B (S12 Figure B for the full range).

**Figure 7:**
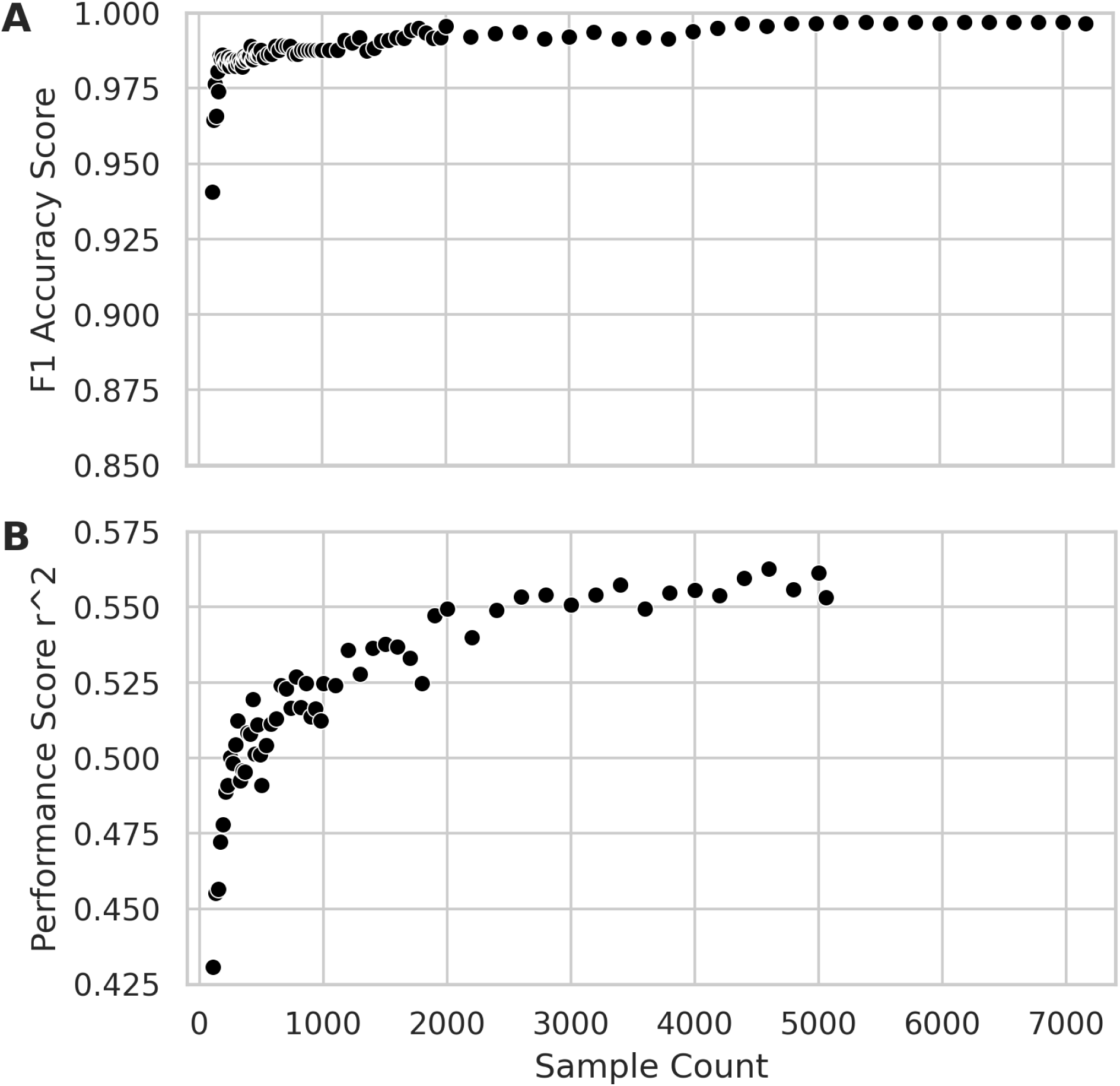
Model performance for different sample counts. Samples were added randomly from the training set, and a new model was created for each sample count. **A)** Model performance for tissue-4 classification dataset **B)** Model performance for DAL regression dataset. Both **A** and **B** are set at a y-axis range of 0.15, which excludes low-count models in preference of readability. For the full range which includes low-count model performance please see **S12 Figure**. Performance is based on testing datasets. Model performance for the tissue-6 dataset is included as **S13 Figure**.

Figure 7: Model performance for different sample counts. Samples were added randomly from the training set, and a new model was created for each sample count. A) Model performance for tissue-4 classification dataset B) Model performance for DAL regression dataset. Both A and B are set at a y-axis range of 0.15, which excludes low-count models in preference of readability. For the full range which includes low-count model performance please see S12 Figure. Performance is based on testing datasets. Model performance for the tissue-6 dataset is included as S13 Figure.

For the randomization approach accounting for BioProject, curves looked similar but were slightly delayed when compared to randomly sampling data (S5 Figure). This is likely due to BioProjects adding only 1 or a few time points. Curves reached the same plateaus as randomly adding samples.

### Boruta and Gene Feature Importance

Boruta feature selection was performed using both the DAL, tissue-4, and tissue-6 datasets to identify genes that were better at predicting their respective labels and age values better than a randomized “shadow” feature of themselves. Boruta creates “shadow” features by randomizing the values of a gene across all samples. These “shadow” features are included during model creation, and if the true feature performs worse than this shadow feature, it is eliminated [22]. For the tissue-6 dataset, 20017 genes were shown to perform better than randomized features (accepted) and 23207 were rejected after 200 runs. For the tissue-4 dataset, 7837 genes were accepted, and 34788 were rejected after 200 runs. For the DAL dataset, 15024 genes were accepted and 26394 were rejected after 200 runs. A reduced GEM was then made for both tissue-6, tissue-4, and DAL by keeping only those genes that were accepted. These reduced datasets represent the genes that perform better than random, however, Boruta does not rank how well they perform. To rank these genes, we used these reduced datasets to perform random forest feature selection. The list of genes and their rankings are available as S6 Table for tissue-6, S7 Table for tissue-4, and S8 Table for the DAL datasets. Figure 8 shows four of the top genes selected as most important for classifying the tissue-4 of the sample. Note the difference in expression between the different samples. Figure 9 shows three of the top genes selected for prediction in the DAL model (after Boruta filtering). Ages falling between days (e.g., 5.5) were rounded up for the sake of plotting. S14 Figure shows 4 of the top genes for the tissue-6 model.

**Figure 8:**
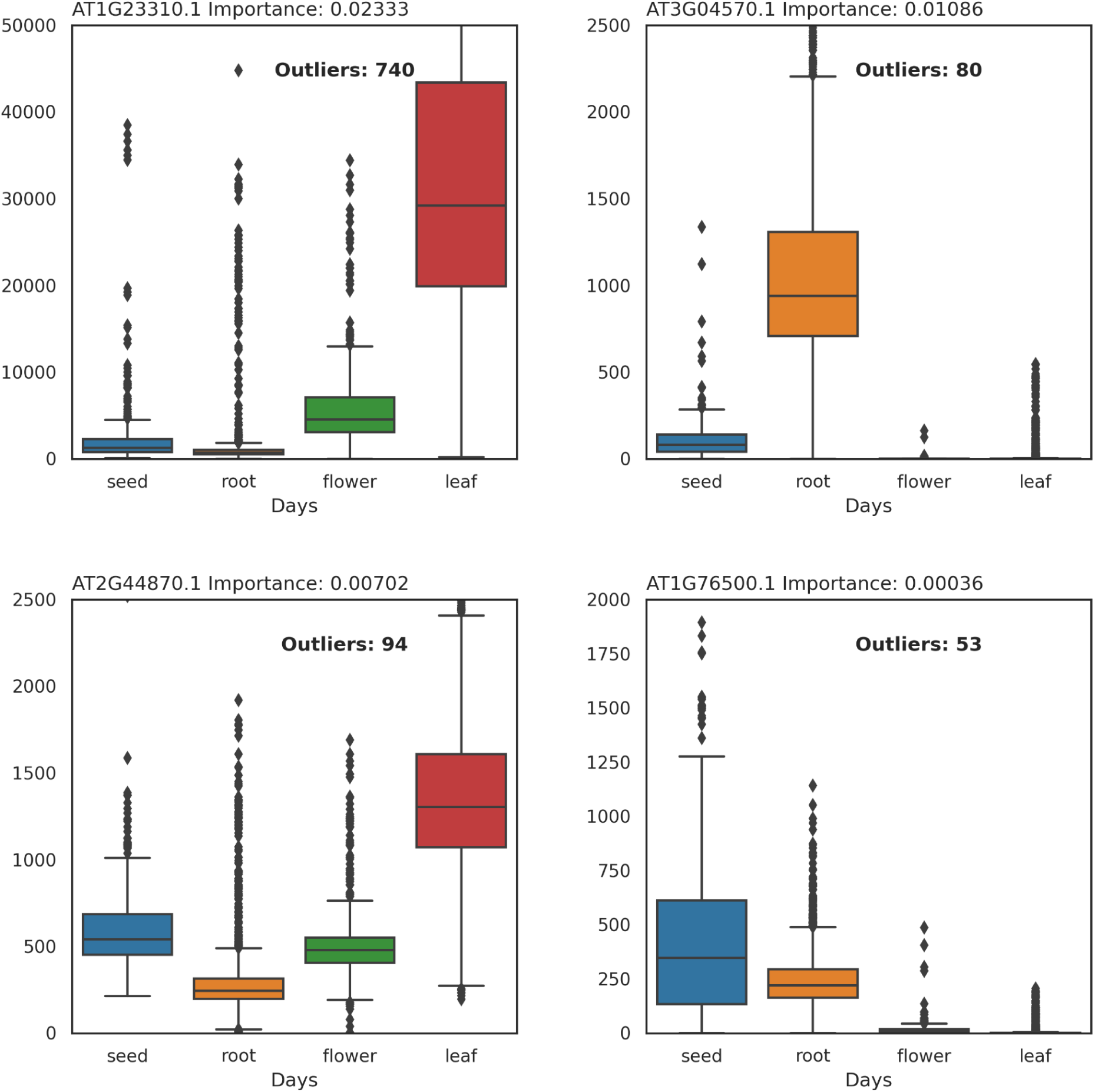
Four of the top-ranked genes from the tissue-4 classification model. Plots show large differences in expression between tissue labels. The number in the panel title next to the gene name is the feature importance of that gene in the model. Figure 8: Four of the top-ranked genes from the tissue-4 classification model. Plots show large differences in expression between tissue labels. The number in the panel title next to the gene name is the feature importance of that gene in the model. Figure 9: Top three genes for the DAL model. The x-axis represents days, and the y-axis represents MRN normalized counts. Ages falling between days (e.g., 5.5) were rounded up for the sake of plotting. For plotting, some outliers were removed to conserve y-axis space.

**Figure 9:**
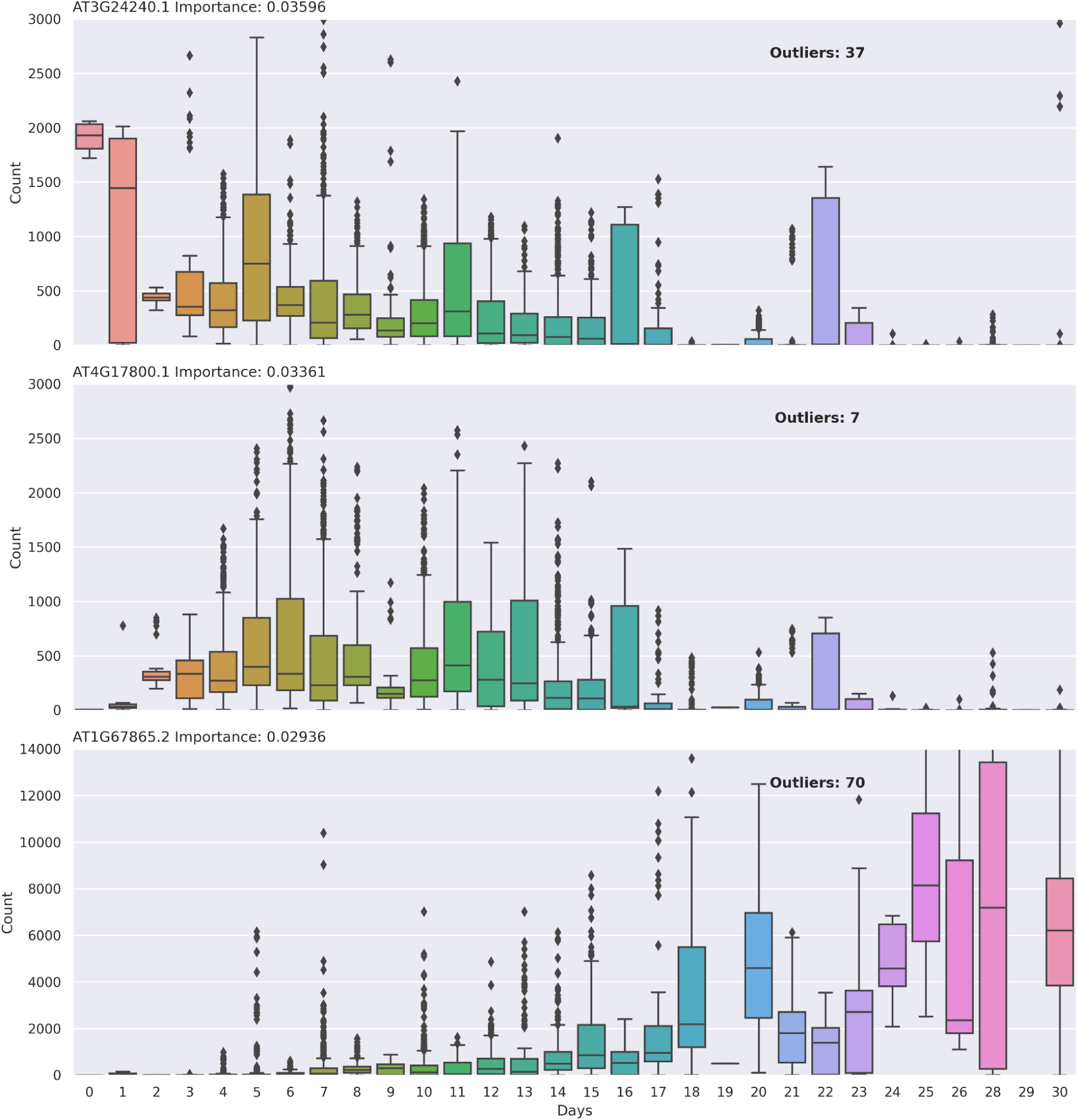
Top three genes for the DAL model. The x-axis represents days, and the y-axis represents MRN normalized counts. Ages falling between days (e.g., 5.5) were rounded up for the sake of plotting. For plotting, some outliers were removed to conserve y-axis space.

## Discussion

For this study, we created a massive Arabidopsis RNA-seq dataset and included metadata such as categorical tissue labels and quantitative age values and used it to explore answers to two questions important for predictive biomarkers traits using gene expression data. These questions are: first, how many samples are required for such models, and second, how do normalization methods affect model performance.

Towards this end, we randomized subsets of the massive Arabidopsis dataset to explore how performance changes as data size increases. We investigated four different normalization approaches. To maximize model performance, we performed random forest parameter selection. To reduce the variables in the data, we applied Boruta and removed genes that were never predictive of the outcome.

### Normalization Method

Four normalization methods were evaluated, TMM, MRN, TPM, and NoNo. In the tissue-6 dataset, the normalization method had no impact on model performance (Figure 2 A). This contrasted with the DAL dataset where a significant increase in model performance was seen for the MRN dataset over the other three methods (Figure 2 B).

MRN normalization takes the geometric mean of each gene across samples and then uses this to calculate a ratio. The median of these ratios taken across each sample is the normalization factor for that sample [35]. This technique considers all samples and all genes in the dataset when calculating these normalization factors. Thus, MRN shares information across samples to identify genes that are not differentially expressed (i.e., stable) and uses those as normalization factors. TMM also does this, but with a weighted mean of log ratios-based strategy [29], whereas TPM does not attempt to identify stable genes and normalizes gene counts in a sample to one million. NoNo data is unnormalized, and differences in sample size can have a major impact on relative expression levels to other datasets. In our experiment, the normalization method did not matter in the classification problem because the genes selected by the model had dramatic differences between tissue types (Figure 8). It seems, therefore, that it does not matter what normalization method is used if genes contributing to a label (such as tissue type) tend to be expressed only in their respective class labels. This contrasted with the age-in-days regression-based models, which relied on continuous, subtle differences between days to create a reliable predictive model (Figure 9). In the age case, if genes are not accurately normalized, the importance of genes can be reduced or not detected. We suspect some causes may be related to sequencing depth and the influence of highly differential genes. We also suspect that the differences in model performance between MRN and TMM normalizations are due to MRN more dramatically changing the underlying data, whereas TMM does not change the counts as much (as illustrated in S2 Figure). Whereas the difference between models may not be very noticeable when normalizing a single dataset [36], normalizing across our massive conglomerate dataset favored the more aggressive normalization of MRN. Further testing will need to be performed to determine if MRN is the best method of normalization for RNA-seq data in other situations.

The differences in how these normalization methods impact the overall gene count of all samples is illustrated in S2 Figure. To summarize, TPM results in all samples having a million counts, TMM results in smoothing when compared to NoNo, and MRN results in more drastic smoothing compared to TMM.

### Accuracy and Model Performance

Accurate phenotype annotations are important for constructing meaningful models. Ensuring accurate phenotypic annotations of samples in large, conglomerate datasets can be difficult. Phenotypic annotations used in this paper came from over 2000 independent scientific experiments, collected by an untold number of researchers. This results in a high likelihood there exist differences in experimental design (e.g., differences in temperature impacting development but not age), chances for errors in reporting (e.g., information entered incorrectly or incorrect samples being uploaded to NCBI), and differences in opinion on how a certain annotation class should be reported (e.g. should age be reported as days after sowing or days after germination?). For conglomerate datasets such as our Arabidopsis data, manual effort was required to reduce the impacts of these annotation irregularities.

The tissue-6, tissue-4, and DAL models were performed with increasing amounts of randomized annotations to assess if the annotations reflected actual biology. The premise was that if the models were overfit and not reflective of true biological variation then we would not see a decrease in model accuracy. However, we saw in all three cases that annotation prediction accuracy decreased. In the DAL models, there was a linear decrease (Figure 6 B) whereas in the tissue-6 (Figure 6 A) and tissue-4 models (S15 Figure) there was a delay in model accuracy decline. This delayed decline in accuracy in the tissue datasets is partially due to the actual amount or randomization being different than the stated amount. This difference is due to the small number of possible tissue types resulting in the same annotation being randomly assigned to a sample (S16 Figure). However, it also shows the robustness of random forest classification models. In these models the true transcriptomic signal needed to accurately predict the testing dataset remained in the model even under modest randomization because the training dataset separated out incorrectly annotated signals into other branches of the forest (what we refer to as “bad branches”. These “bad branches” accurately predicted poorly annotated labels but did not impact the good branches needed to predict the testing dataset, allowing for a high training accuracy.

This highlights an advantage of random forest classification that it can be fairly robust to poorly annotated data if the majority of data is accurately classified and there is true biological variation present. Random forest regression is also recalcitrant to outliers and poorly annotated data, but because it is predicting a continuous variable via averages it shows a linear decrease in performance with increasing randomness.

The two tissue models, tissue-6 and tissue-4, had excellent accuracy. However, tissue-6 illustrates the issue in large conglomerate datasets of annotation specificity. Of the 240 samples in the testing set labeled as “shoot” only 8 of them were accurately labeled as such by our prediction model (3.3% accuracy) (Figure 3 B and D). The vast majority of “shoot” labeled samples were predicated as either “seedling” or “leaf”. This is likely because “shoot” is an ambiguous label when considering Arabidopsis plants.

Whereas a young seedling (stage 0-1) [19] has an obvious shoot, an older Arabidopsis plant does not. This results in the shoot label sharing a large amount of biological similarity with either the seedling label or the leaf label (which includes full rosette) making them difficult to distinguish. The other poorly defined term is “seedling”, which also has subjectivity built into it. There is not a well-defined annotation for what an “Arabidopsis seedling” is, so we decided to define it as any plant before “stage one” which is around 6 days post-germination [19]. However, many of the tissue annotations were sparse and we were not able to distinguish if a “seedling” annotation fit our criteria. It is likely that a large amount of “seedling” labeled samples do not fit our definition, which is illustrated by some of the available annotations being, “leaf (20-day seedling)”, “15-day-old seedling” and “21 day old seedling”. These should be plants that are well beyond this stage, and in some instances with several sets of true leaves. The reduced tissue-4 model mitigates these issues by removing the ambiguous labels, and instead concentrates solely on four well-defined categories, resulting in excellent accuracy. However, it trades this increased accuracy for a reduced number of annotation categories.

The DAL age dataset was also subject to inaccuracies in labeling. This dataset consisted of only samples which were annotated with the word “day” or equivalent. It is unclear if the meaning of each of these labels is referring to “days after sowing”, “days after germination” or “days after stratification”. It could be that these dates should be shifted anywhere from 0-6 days if they are to represent the same timescale. This uncertainty in actual date precision is captured in our DAL model, with the testing dataset having an RMSE of 4.4423, which can be thought of as the average number of days the prediction deviates from the true age. The DAL model was able to capture ∼55% of the variability in the DAL dataset, which is good considering the issues with annotations.

To explore if we could mitigate the issue of precision with the age annotations, we tested two additional datasets: DAG and DAS. The DAG and DAS age categories were more precise in how age was reported on NCBI, specifying whether the measurements are referring to time after germination or sowing respectively.

Unfortunately, the more specific annotations for DAG and DAS failed to achieve higher model accuracy when compared to DAL (S9 Figure). This is because we did not have as many samples from as many BioProjects to create models. However, if we compare the performance of the DAG dataset (r^2^ = .45, S9 Figure A, B, C) to that of the DAL model created using a comparable number of BioProjects (45 BioProjects)(r^2^ = .28, S4 Figure 4 C) we see better performance of the DAG dataset than the DAL. This illustrates how improved precision in age annotations can potentially make better models. We encourage researchers to submit as accurate information as possible when submitting BioSample data about their projects to NCBI. It is useful to consider FAIR (Findable, Accessible, Interoperable, Reusable) guiding principles when submitting datasets [37].

### Number of Samples Required

Two of the main goals of this project were to assess how many RNA-seq samples are required for model creation and if samples from disparate experiments could be used together. These goals were explored for both the categorical tissue datasets (tissue-6 and tissue-4) as well as the quantitative age dataset (DAL). Tissue classification rapidly achieved high performance after only a few hundred samples were added, whereas age prediction performance only reached a plateau after 2-3 thousand samples (Figure 7 A and B). This is likely due to the differences in the complexity of the models. The tissue classification problem was only attempting to classify samples into 6 or 4 categories, with each of these being distinct tissues of Arabidopsis. It is known that there are a number of genes that are only active in certain tissues [38]. In contrast, the age regression model was over a continuous range of 0-30 days. In addition to the large number of ages of the sample (0-30 days) the model also had to consider all the other latent variables present within these samples. These latent variables include differences in genotype, temperature, moisture level, nutrient availability, gene knockouts, growth media, pest pressure, and tissue type. All these factors may have an impact on gene expression in a manner semi-independent of gene expression related to age. This means that age models must either identify genes unimpacted by these latent variables or independently predict age by considering these latent variables. In reality, it is most likely a trade-off between these two cases, where the model includes genes mostly predictive of only age and genes that take into account other variables that may impact the age variable (Figure 9). While tissue-related expression patterns are also impacted by these latent variables, it appears that tissue type differentiation has unique enough expression patterns to accurately differentiate between them (tissue-4 Figure 8, tissue-6 S14 Figure).

Supplementing the DAL dataset using SMOTE resulted in a modest increase in annotation accuracy. SMOTE is intended to be used on classification datasets, not continuous datasets, so to use SMOTE on the DAL dataset we rounded the days and converted them to distinct categories. After converting back to numerical values, we saw a modest increase in model performance (S11 Figure). This increased performance, because of balancing by SMOTE, illustrates the importance of balanced datasets for model prediction. Balancing unbalanced datasets is a currently active area of research, especially for continuous variable models [39], and a consensus on how to balance tabular datasets, such as ours, does not have an agreed-upon solution. Here we show that using SMOTE on our continuous variable age had a modest increase in model performance. Ultimately, this increase was not dramatic enough to warrant the decrease in model interpretability resulting from introducing artificially generated data.

Here we do not propose a definitive cutoff for the number of samples required to create an accurate predictive model using RNA-seq datasets. Differences in datasets, experimental design, biological impact, and sample annotation quality will have an impact on model accuracy, so broad recommendations are warranted. As already stated, tissues tend to have genes that are uniquely expressed thus this variable represents a “simple” model, or one with fewer multi-functional, co-dependent, or conditional relationships amongst genes. Thus, we expect the tissue model to represent a lower bound in terms of the number of required samples. We point readers to Figure 7 and suggest that biomarker experiments should include at least a few hundred samples for accurate prediction. The age models indicate that for more complicated traits (such as for environmentally impacted traits), a few thousand samples may be required.

## Conclusion

The research presented here provides foundational knowledge about possible sample size requirements and the effects of RNA-seq normalization for transcriptomic biomarker model development. Results provide guidance on the minimum number of samples and normalization methods that may be needed for accurate models. Such guidance has applications for the development of predictive transcriptomic biomarkers which are gaining popularity in precision medicine and specialty agricultural crops.

Finally, we encourage researchers submitting RNA-seq samples to NCBI, or other public repositories, to provide correct and comparable annotations for their samples based on FAIR data principles [37].

## Author’s Contributions

**JAH** Conceptualization, Methodology, Software, Formal Analysis, Investigation, Data Curation, Writing – Original Draft, Writing – Review & Editing, Visualization, Project administration. **LAH** Funding acquisition, Writing – Review & Editing. **SPF** Funding acquisition, Supervision, Project administration, Writing – Review & Editing.

## Data Availability Statement

All normalized gene expression datasets, phenotype datasets, and intermediary files created for this research are publically available on Zenodo at link https://zenodo.org/doi/10.5281/zenodo.10183150

All code written in support of this publication is publicly available on GitLab at link https://gitlab.com/ficklinlab-public/modeling-with-transcriptomics

## List of Abbreviations

Arabidopsis: *Arabidopsis thaliana*
DAG: dataset containing samples labeled as Days After Germination
DAL: dataset containing samples “Days” Age annotation Labeled
DAS: dataset containing samples labeled as Days After Sowing
GEM: Gene Expression Matrix
HSD: Tukey’s Honestly Significant Difference
MRN: Median Ratios Normalization
NCBI: National Center for Biotechnology Information
NoNo: No Normalization
SMOTE: Synthetic Minority Over-sampling Technique
SRA: Sequence Read Archive
TAIR: The Arabidopsis Information Resource
tissue-4: dataset containing samples annotated as “leaf”, “seed”, “root”, and “flower”
tissue-6: dataset containing samples annotated as “leaf”, “seedling”, “shoot”, “seed”, “root”, and “flower”
TMM: Trimmed Mean of M values
TPM: Transcripts Per kilobase Million

## Supplemental Materials

### Tables

**S1 Table. Summary of the splits for the Tissue Dataset.** Shows total sample counts and number of BioProjects in each. Most BioProjects consisted of a single type of tissue type, but a few consisted of multiple which is why a sum of the BioProjects Train does not match with total number of BioProjects.

**S2 Table. Tukey HSD between the different normalization methods for Age.** MRN was significantly higher than the other normalization methods.

**S3 Table. Parameter Optimization Results for RandomForest Classification Tissue.** Optimal parameters were ‘n_estimators’: 1500, ‘min_samples_leaf’: 3, ‘max_features’: ‘sqrt’, ‘max_depth’: 48, ‘bootstrap’: False.

**S4 Table. Parameter Optimization Results for RandomForest Regression Age.** Optimal parameters were *’n_estimators’*: 1500*, ‘min_samples_leaf’*: 3, *’max_features’*: ‘sqrt’, *’max_depth’*: 48, *’bootstrap’*: False

**S5 Table. Predictions for 32432 RNA-seq Samples**. Prediction columns for the three models are: ‘tissue_6_prediction’, ‘tissue_4_prediction’,’DAL_prediction’. Ground truth annotation columns are ‘tissue’, ‘days_age’, ‘age_category’, and ‘age_category_full_name’. Note that ground truth columns do not have values for every RNA-seq sample, as all samples do not have ground truth information about their annotations. The first three columns are information about the RNA-seq sample name: ‘experiment’, BioProject: ‘bioproject_name’, and BioSample: ‘biosample_name’. Also has information about if the sample was in the training or testing set for each model: ‘tissue_4_train_test’, ‘tissue_6_train_test’, ‘DAL_train_test’.

**S6 Table. Feature Importance Tissue-6 After Boruta.** List of genes and their importance in the Tissue-6 model.

**S7 Table. Feature Importance Tissue-4 After Boruta.** List of genes and their importance for the Tissue-4 model.

**S8 Table. Feature Importance DAL After Boruta.** List of genes and their importance for the DAL model.

### Supplemental Figures

**S1 Figure. Number of features remaining for different thresholds.** Number of features remaining that are over m count for n samples (x-axis). **A)** Number of genes remaining for TPM with different minimum counts (m), **B)** Same except for NoNo. Red Line is at 1000 samples. Final thresholding was based on the NoNo dataset, with all other datasets (MRN, TMM, and TPM) containing the same genes.

**S2 Figure. Histogram of total sample read count for the four normalizations.** The x-axis is log2 transformed and is the same for each plot. The y-axis maximum value is different in each plot. *Abbreviations:* Trimmed Mean of M values (TMM), Median of Ratios Normalization (MRN), Transcripts per kilobase million (TPM), unnormalized count data (NoNo).

**S3 Figure. Phenotype annotation class.** Diagram showing the sparsity of data. The annotations retrieved are very sparse, with many annotation columns only used for one or a few BioProjects. The annotations (Tissue and Age) we use for this project are highlighted in color. The x-axis represents the number of BioProjects which use an annotation, and the y-axis represents the number of annotations which are at that category. We were able to highlight tissue and age because they are the only annotation which is present for that number of BioProjects.

**S4 Figure. Age categories by timepoint.** Annotations for age were split into categories based on how the BioSample information for age was reported (i.e. if sample age information was reported as “10 Days after Germination” It would be reported as “Days after Germination (DAG)” whereas if it was just a number “10”, then it would be reported as “Number Only”). Categories were: ““Days” Age Annotation Labeled (DAL)”, “Weeks”, “Months”, “Number Only”, “Days After Sowing (DAS)”, “Days After Germination (DAG)”, “Days After Pollination”, and “Days After Flowering”. Additional information can be found in Table 2 in the text.

**S5 Figure. Model Performance Adding by BioProjects.** BioProjects were added 10 at a time. Performance is always measured on the respective testing dataset. **A)** Tissue-6 Performance. **B)** Tissue-4 performance **C)** DAL Performance. The x-axis are the same for each model.

**S6 Figure. SMOTE Resampling of Regression Data.** Dates with half days were rounded up to the nearest date. Dates with sample count under 10 samples were removed before running SMOTE (0,1,19,27, 29). **A)** Sample distribution before SMOTE **B)** Sample distribution after SMOTE

**S7 Figure. GridSearchCV of different *max_feature* depths.** GridSearchCV of different max_feature depths (x-axis) for the **A)** tissue-6 classification model and **B)** DAL regression model. Note that y-axis is scaled at 0.025 between ticks for both plots.

**S8 Figure. Coloration of** Figure 2 **as “max_depth”.** Coloration of Figure 2 as “max_depth” to illustrate that the very low points are just too low of “max_depth”

**S9 Figure. Age models DAG and DAS.** Age models using samples classified as Days After Germination (DAG)(**A,B,C**) and Days After Sowing (DAS)(**D,E,F**). Plots **A** and **B** represent the distribution of samples for DAG and DAS respectively. Plots **B** nd **C** represent the training and testing performance (r^2^) for DAG and **E** and **F** represent the training and testing performance for DAS. DAG model was created using 574 samples across 52 BioProjects, and the DAS model was created using 804 samples across 19 BioProjects.

**S10 Figure. DAG and DAS performance on DAL dataset.** DAG model (A and B) and the DAS model (C and D) performance on the DAL dataset. DAL dataset is split between the same train and test splits used for the DAL model. DAG and DAS model performance was lower than the DAL model on the DAL dataset.

**S11 Figure. SMOTE DAL dataset model performance. A** is training performance, and **B** is testing performance.

**S12 Figure. Full range model performance tissue-4 and DAL.** Full range model performance for different sample counts. This is the same figure as Figure 7 except the entire y-axis is shown. **A)** Model performance for tissue-4 classification dataset **B)** Model performance for DAL regression dataset.

**S13 Figure. Model Performance of tissue-6 dataset randomly added.** This is complementary to Figure 7 which shows tissue-4 and DAL datasets.

**S14 Figure. Four of the top genes (features) from the tissue-6 dataset.** X-axis is divides samples by tissue. Plots y-axis are scaled according to expression level of each gene (y-axis) with the number of outliers not appearing on the plot reported in the top right corner.

**S15 Figure. Accuracy results for tissue-4 randomization.** Reported for Train (solid lines) and Test (dashed lines) for both Accuracy (black lines) and F1 (red lines).

**S16 Figure. Actual Random Plot for DAL, tissue-4 and tissue-6 datasets.** Due to tissue-4 and tissue-6 having only 4 and 6 categories respectively, during randomization, there is still a relatively large chance that the same label is assigned. The x-axis represents what percent of the data is randomized, and the y-axis shows the actual percent, once randomly assigning the same variable to self is taken into account. Notice that DAL is more or less a 45-degree line, whereas tissue-6 and tissue-4 have less actual percent randomized.https://gitlab.com/ficklinlab-public/modeling-with-transcriptomics

